# Identification of treatment-responsive genes in spatial transcriptomics data by leveraging injection site information

**DOI:** 10.1101/2023.06.30.547203

**Authors:** Felicita Pia Masone, Francesco Napolitano

## Abstract

Spatial Transcriptomics assays allow to study gene expression as a function of the spatial position of cells across a tissue sample. Although several methods have been proposed to identify spatially variable genes, they do not take into account the position of the injection site in the case of treated samples. In this study, we developed a method to identify treatment-responsive genes based on the assumption that the distance of the cells from the injection site across the tissue would affect the corresponding transcriptional response. In particular, we tested our approach using a publicly available ST dataset obtained after injection of heme into the striatum nucleus of a murine brain. We observed that several biologically relevant genes were detected by our method as showing a distance-dependent expression trend. We finally compared the results against a ground-truth gene set and a state-of art pattern-based method.

## 1 Introduction

Spatial transcriptomics (ST) is a relatively new technology that enables spatially resolved analysis of gene expression within a tissue or cell at the whole transcriptome level. ST has the potential to provide detailed information about the biology, organization, and intercellular communication of cells and tissues, and it can be used in a wide range of biomedical applications, including the study of disease, cell development and differentiation, immunology, neuroscience, and pharmacology [1]. A typical application of non-spatial transcriptomic assays (such as RNAseq) is differential expression (DE) analysis, in which different biological conditions are compared against each other with the aim of identifying condition-dependent expression. One such example is the analysis of treated versus non-treated cells. On the other hand, ST data analysis typically explores internal differences among spatial regions in a tissue. For example, it is possible to search for genes that exhibit spatial expression patterns, called spatially variable genes (SVGs, [2]). In the case of differential analysis, ST datasets are typically analyzed separately to extract relevant genes from each condition. The different results are then compared [3]. Moreover, potentially useful information about the injection site for a treatment, which can be technically exploited in an ST context, is usually ignored.

In this paper, we propose an alternative method for the identification of SVGs in samples subjected to injection of a foreign substance called RGDIST (Responsive Genes based on Distance from Injection SiTe). Our approach is based on the assumption that treatment-responsive genes will show an expression alteration that is affected by the distance of each cell from the injection site. Moreover, in order to reduce tissue-specific transcriptional effects that are not driven by the treatment, we use batch effect removal based on cell types. This allows for the identification of several “differentially expressed” genes even in the absence of a reference condition, such as an untreated sample. In order to demonstrate the efficacy of our approach, we compared it with a baseline expression model only using average gene expression, a state-of-art SVG detection algorithm (spatialDE), and an ad hoc treatment-control differential analysis. We used a previously published ST dataset [3] obtained after injection of heme into the striatal nucleus of a mouse brain.

## 2 Data and Methods

The ST data used for our analysis are publicly available from the Gene Expression Omnibus website (accession number GSE182127). The dataset was obtained after stereotactic injections of heme and hemopexin into a mouse brain. The rationale for the experiment is that many brain diseases such as stroke, chronic hypertension, amyloid angiopathy, vascular malformations and brain tumors can cause intracranial hemorrhage [4], after which heme is released from cell-free hemoglobin and can cause secondary brain injury in the bleeding area. The study by Buzzi et al. aimed to spatially analyze gene expression changes in the mouse brain after heme injection thus simulating intracranial hemorrhage. The dataset includes seven samples of coronal sections: two untreated controls, four treated with increasing concentrations of heme and one treated with heme-hemopexin. We chose to use the sample treated with 10 nmol of heme (GSM5519059) as the one likely showing the most pronounced transcriptional effects. We performed our analysis within the R environment (SpatialExperiment package v1.8.1 [5]). To preprocess the data, we followed a standard analysis pipeline for scRNAseq data that is also used for ST data [6]. The dataset includes coordinates of the injection site and manually assigned anatomical clusters. We excluded from further analysis: (i) spots not assigned to any anatomical cluster, as this information is required in subsequent steps of our analysis; (ii) genes detected in less than three spots, since at least three observations are required to subsequently compute correlations. Then we computed the Euclidean distance of each spot from the injection site. Since the injection site is provided as an area, we chose its central spot to compute the distances. In this instance, the injection area encompasses 158 spots out of 2,310 (6.84% of the total spot). In order to remove tissue-driven transcriptional effects that are not dependent from the treatment, we performed a batch correction (BC) of the expression data using the ComBat seq approach (R package sva v3.35.2) [7] in which we considered the anatomical clusters as batches. We then normalized the corrected counts by their total number of transcripts within each spot.

To identify genes affected by the treatment, we computed the Spearman’s rank correlation coefficient between the expression of each gene across all the spots and the corresponding distance from each spot to the injection site, excluding null counts. The Spearman coefficient was chosen as the main objective of the analysis is the identification of monotonic trends. We performed the same analyses both on non-corrected and batch-corrected expression data to evaluate the effects of the correction step. In order to extract a set of genes responding to treatment, we used a p-value threshold of 0.01 and a correlation threshold of ±0.5 for the data before correction. The same selection approach was applied to the BC data, but the correlation threshold was set at ±0.35. Such thresholds were chosen on the basis of a volcano-plot like analysis in which differential expression was replaced by the correlation between expression and distances (see Figure 1). To demonstrate the validity of RGDIST, we ranked all the genes according to the Benjamini-Hochberg corrected Spearman Correlation p-values and then compared such ranking with those obtained by two other methods using different levels of spatial information. First, we devised a baseline method solely based on gene expression without neither control nor spatial information. In this case, we just used the average expression of each gene across the tissue sample, normalized by the average expression of all the genes. Then, we applied a state-of-art method for the identification of SVGs named SpatialDE [2] which uses spatial information from the treated sample only. We applied SpatialDE following its standard pipeline (R package spatialDE v1.4.3) and ranked the genes according to their Log Likelihood Ratio (LLR) values in descending order. Finally, we used the set of significantly up- and down-regulated genes in response to heme treatment as identified by the authors of the ST dataset [3]. These genes were selected by pooling the results of two different approaches, which included both spatial and treatment-control differential analysis, together with an anatomy-based and a clustering-based cell type identification. The final set includes a total of 62 up-regulated genes and 6 down-regulated genes. We used such genes as a ground-truth to evaluate all the methods against and then considered the meaning of the resulting discrepancies. Tissue expression pattern visualizations were obtained using the R package `spatialLIBD` v1.10.1 [8].

**Figure 1:**
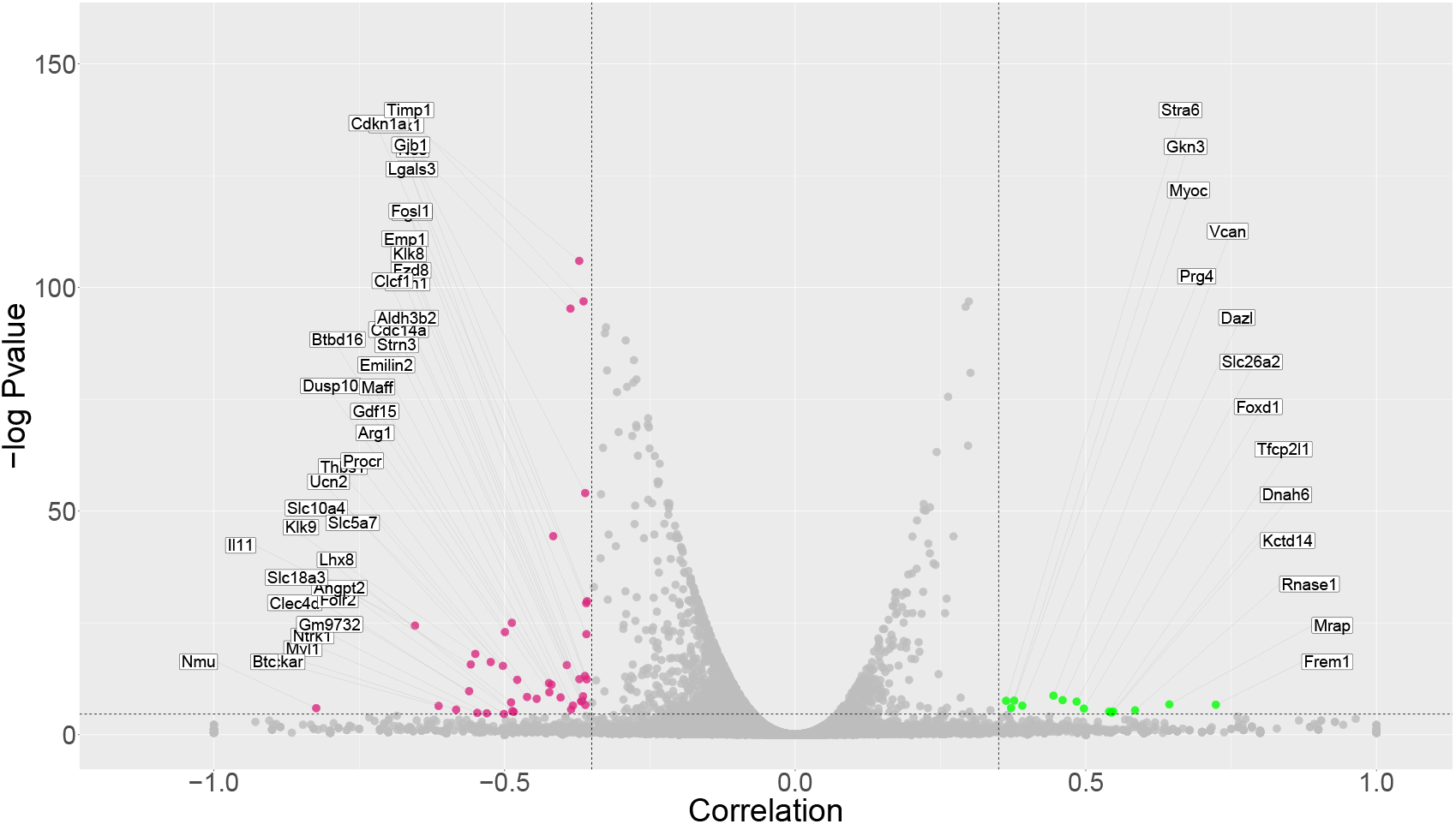
Selection of significant genes after Batch Correction was performed by setting a threshold of 0.01 for p-values and ±0.35 for correlation values. Correlated genes are shown in green (right), anticorrelated genes are shown in pink (left).

## 3 Results

Our analysis revealed two groups of significant genes (Figure 1): correlated (9 before BC; 14 after BC), which show an upward trend in their expression as the distance from the injection site increases; and anticorrelated (28 before BC; 42 after BC), which show a downward trend. The spatialDE method detected 4,280 genes that exhibit a spatial pattern, including all our significant genes, although they were ranked low in comparison. Results before BC are not shown.

Among our significant genes, some of particular interest are reported in Table 1. Their expression patterns over the tissue and their expression values as a function of distance from the injection site are visualized in the Figure 2. These genes are not part of the ground truth set and are all ranked low by spatialDE.

**Table 1:** Significant genes of particular interest ranked by correlation p-values

| Gene | Correlation | Pvalue | RGDIST rank | SpatialDE rank | Comment |
| --- | --- | --- | --- | --- | --- |
| Il33 | -0.37 | $9.7 \times 10^{-47}$ | 1 | 215 | Anti-inflammatory cytokine [9] |
| Il11 | -0.65 | $2.6 \times 10^{-11}$ | 9 | 2024 | Anti-inflammatory cytokine [10] |
| Gdf15 | -0.42 | $9.7 \times 10^{-06}$ | 21 | 494 | Inflammatory marker [11] |
| Procr | -0.44 | $3.3 \times 10^{-04}$ | 28 | 1391 | Triggers cytoprotective pathway [12] |
| Aldh3b2 | -0.37 | $5.9 \times 10^{-04}$ | 30 | 3235 | Removes ROS-derived aldehyde [13] |

**Figure 2:**
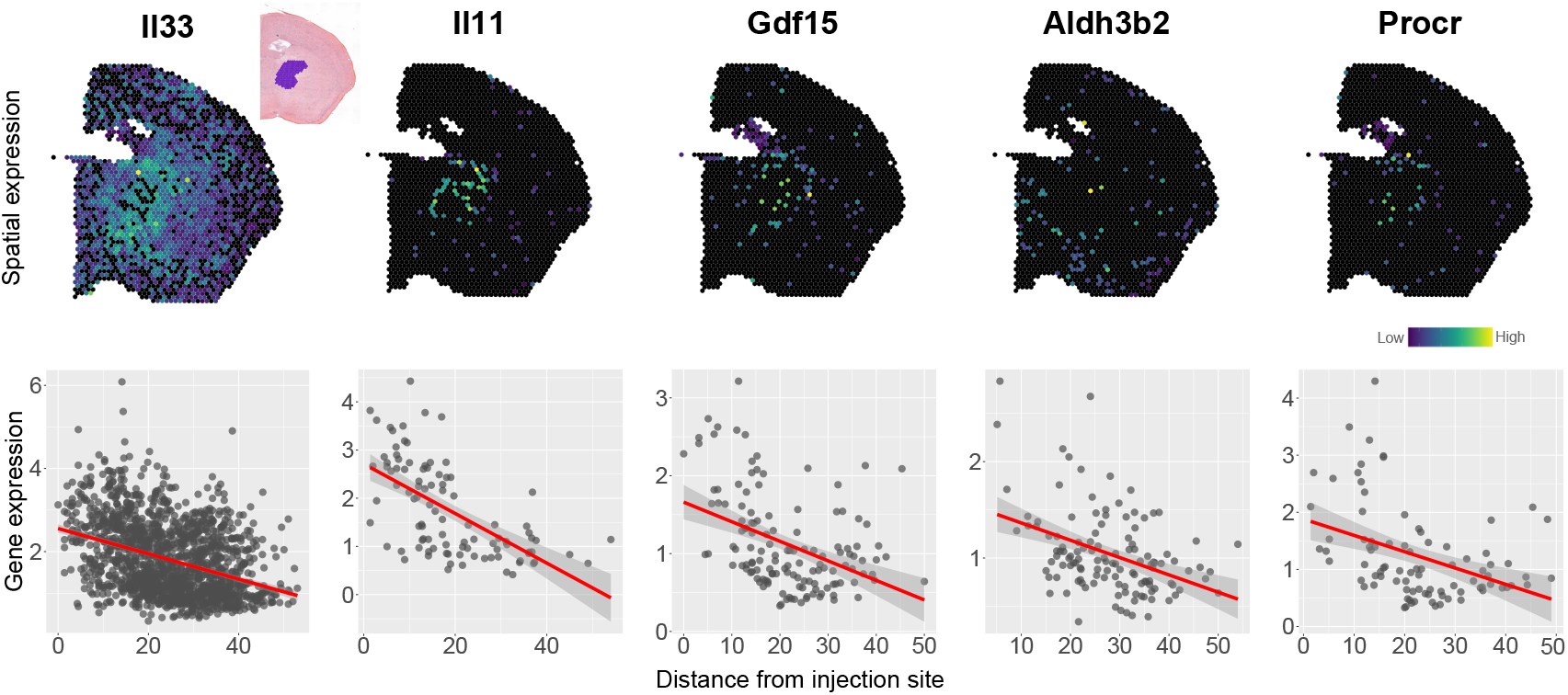
First row: spatial visualization of selected genes expression. Inset in the first panel shows injection site area. Second row: distance-expression plot of selected genes.

Concerning such genes, IL-33 protein is a tissue-derived nuclear cytokine from the IL-1 family. It is expressed both during homeostasis and inflammation and acts as an alarm signal released upon cell injury or tissue damage to alert immune cells. In mice, after brain injury post-mitotic OLIG2+ oligodendrocytes rapidly release IL-33 to promote recovery [9]. IL-11 is a member of the IL-6 family of cytokines and has been demonstrated to have various immunological effects on both innate and adaptive immune processes. *In vitro* studies have shown that IL-11 functions as an anti-inflammatory cytokine by modifying the effector function of macrophages [10]. The Gdf15 gene encodes Growth and Differentiation Factor 15 (GDF15), a member of the transforming growth factor-*β* (TGF-*β*) superfamily of proteins. GDF15 functions as an inflammatory marker and plays a critical role in regulating endothelial adaptations after vascular damage. Data supports the potential use of GDF15 as a biomarker in stroke patients [11]. Another example is the Procr gene, which encodes the endothelial protein C receptor (PROCR). It has been showed that PROCR activates intracellular signaling pathways, leading to cytoprotective effects. When Activated Protein C (APC) binds to cells expressing PROCR, it triggers multiple beneficial cytoprotective activities that suppress apoptosis, inflammation, and breakdown of the endothelial barrier [12]. The Aldh3b2 gene encodes for a member of Aldehyde dehydrogenase 3 family, ALDH3B2. This enzyme involved in metabolism of ROS-generated aldehyde products and it shows a wide substrate preference from medium-chain to long-chain aldehydes. It has been observed that ALDH3B2 is localized to lipid droplets [13], which are frequently observed in cultured neurons and their number is increased in response to cellular stress, e.g. due to a stroke [14].

In order to investigate cellular mechanisms involving the genes identified by RGDIST, we conducted a gene set enrichment analysis using the DAVID online platform [15]. The resulting enriched gene sets involved a variety of relevant mechanisms, including *responses to xenobiotic stimuli* (*p ≈* 2.9× 10^*−*4^), *cytokine activity* (*p ≈* 1.3 ×10^*−*3^), *cell death* (*p ≈* 2.5 ×10^*−*3^), *growth factor activity* (*p ≈* 3.6 ×10^*−*3^), *positive regulation of programmed cell death* (*p ≈* 2.1 ×10^*−*2^), *positive regulation of reactive oxygen species metabolic process* (*p ≈* 6.1 ×10^*−*2^), *response to axon injury* (*p ≈* 7.3 ×10^*−*2^). These results are consistent with the effects caused by the presence of free hemoglobin in tissues. Analogous pathways emerge from analyzing ground-truth genes and genes identified by spatialDEs (data not shown). Free heme primarily induces the formation of ROS, which can cause oxidative damage to the surrounding environment and trigger immune responses by the body. Additionally, free heme can interact with cytoplasmic receptors and activate intracellular signaling pathways [3].

The boxplot in Figure 3a compares the three methods showing that both RGDIST and spatialDE are accurate in identifying significant genes in response to heme without using a control sample. SpatialDE tends to rank ground-truth genes high (median: 537.5) based on spatial patterns in their expression. On the other hand, RGDIST ranks them even higher overall (median: 391), although several genes are ranked significantly lower as they do not exhibit a trend in expression (see Figure 3b). Furthermore, Figure 3c displays plots for two significant genes for the spatialDE method that exhibit a neat spatial pattern (Myl4, rank: 45; Tac1, rank: 85). Considering the expression patterns across the tissue sample and the lack of a trend with respect to the injection site, we hypothesize that the expression of such genes, which is deemed relevant by other methods, may be affected by residual cell-type dependent effects convoluted with the heme treatment.

**Figure 3:**
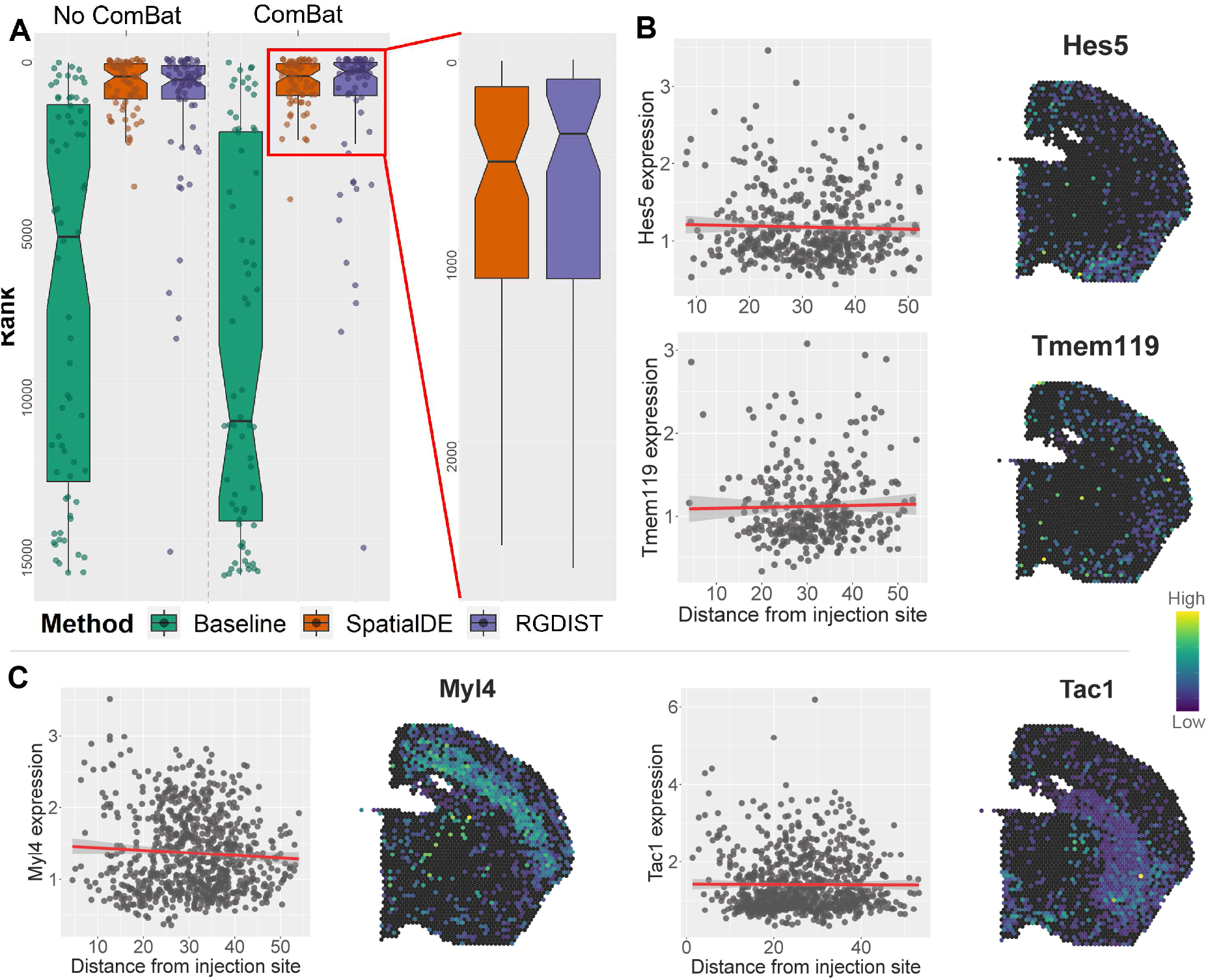
a) Performance of the different methods with and without ComBat in ranking the ground-truth genes. Best performing methods are zoomed in on the right. b) Spatial visualization and distance-expression plots of two ground truth genes that are not deemed significant by RGDIST. c) Spatial visualization and distance-expression plots of two genes deemed significant by spatialDE and not by RGDIST.

## 4 Conclusion

RGDIST was demonstrated to be an effective tool in identifying treatment responsive genes in TS data without using data from the control sample. Future studies are needed to improve the method. For example, its sensitivity to the accuracy of the injection site position or to high treatment times or doses. Moreover, RGDIST could be used to infer the injection site based on the correlations computed at each spot. All such points will be the main aims of further investigations.

